# Cumulative Transfer Function for Assessment of MRI-Induced RF Heating Risk in Pediatric Patients Implanted with Bifurcated Leads

**DOI:** 10.64898/2026.07.08.737115

**Authors:** Fuchang Jiang, Jasmine Vu, Bhumi Bhusal, Yifeng Qian, Safa Hameed, Daniel Kim, Gregory Webster, Giorgio Bonmassar, Laleh Golestanirad

## Abstract

**Purpose:** RF-induced heating remains a major barrier to MRI access for patients with epicardial cardiac implantable electronic devices (CIEDs). Although ISO/TS 10974 Tier-3 transfer function (TF) methods are established for unbranched leads, no analogous framework exists for bifurcated leads, in which branch asymmetry and inter-branch coupling may substantially alter heating. We developed and validated a cumulative transfer function (cTF) framework to address this gap.

**Methods:** Following ISO/TS 10974 Tier-3 formalism, we measured, calibrated, and validated cTFs for a commercial 35 cm bipolar epicardial lead at 1.5 T. The framework explicitly accounts for branch-specific response and cross-branch coupling. Validation was performed with 24 canonical lead configurations in a homogeneous phantom and, without recalibration, in a heterogeneous anthropomorphic pediatric phantom with clinically derived trajectories. A single-branch TF approximation served as a comparator. The validated cTF was applied to predict RF heating across adult and pediatric human models at multiple imaging landmarks.

**Results:** Compared with the single-branch TF approximation, the cTF reduced prediction error by nearly 70% in the primary validation dataset. In secondary validation, the cTF maintained low error across clinically relevant trajectories and imaging landmarks. In human models, the framework revealed marked anatomy- and landmark-dependent variation in predicted heating for the tested 35 cm lead, with low predicted heating in pediatric models and substantially higher heating in selected adult chest and upper abdominal imaging scenarios.

**Conclusion:** The cTF provides a validated framework for RF-heating assessment of bifurcated leads and substantially improves prediction accuracy over single-branch TF approximations that neglect branch coupling.

## 1. Introduction

Children with congenital heart disease are increasingly living for decades with epicardial cardiac implantable electronic devices (CIEDs). As survival improves and indications expand, many of these patients require longitudinal imaging over the course of childhood and adulthood.^1-3^ MRI is particularly valuable in this setting because it provides high soft-tissue contrast without ionizing radiation, yet access remains limited for patients with epicardial systems.^4, 5^ This limitation has important consequences, as children with CIEDs may be diverted to alternative imaging modalities and have been reported to receive a fourfold higher cumulative radiation dose than age- and disease-matched peers without CIEDs.^6, 7^ Notably, an expert review estimated that ∼80% of those CT examinations would have been performed with MRI if heating risk were established.^6^

A major obstacle is that epicardial systems commonly use bifurcated leads, in which a single lead body splits into two terminal branches connected to separate electrodes. This geometry differs fundamentally from the unbranched leads for which current MRI RF-heating assessment frameworks were developed. In bifurcated leads, the induced current at one electrode may depend not only on the local incident electric field along its own branch, but also on coupling through the second branch and shared trunk. As a result, RF-heating behavior cannot, in general, be captured by a single conventional transfer function (TF).

The current gold standard for quantifying MRI-induced RF heating near implanted leads is the Tier-3 TF approach outlined in ISO/TS 10974.^8^ Introduced by Park et al.^9^ and refined by subsequent work,^10-15^ the TF framework was developed for simple, unbranched lead geometries (Fig. 1a). It therefore does not capture current division at a bifurcation or electromagnetic coupling between the branches of a bifurcated lead (Fig 1b). Growing clinical use of multi-channel systems and multiple devices has intensified interest in coupling phenomena; prior studies have examined parallel, unconnected leads routed in close proximity (Fig. 1c).^16, 17^ However, a TF methodology tailored to bifurcated structures—one that explicitly accounts for junction and inter-branch interactions— remains an unmet need.

**Figure 1:**
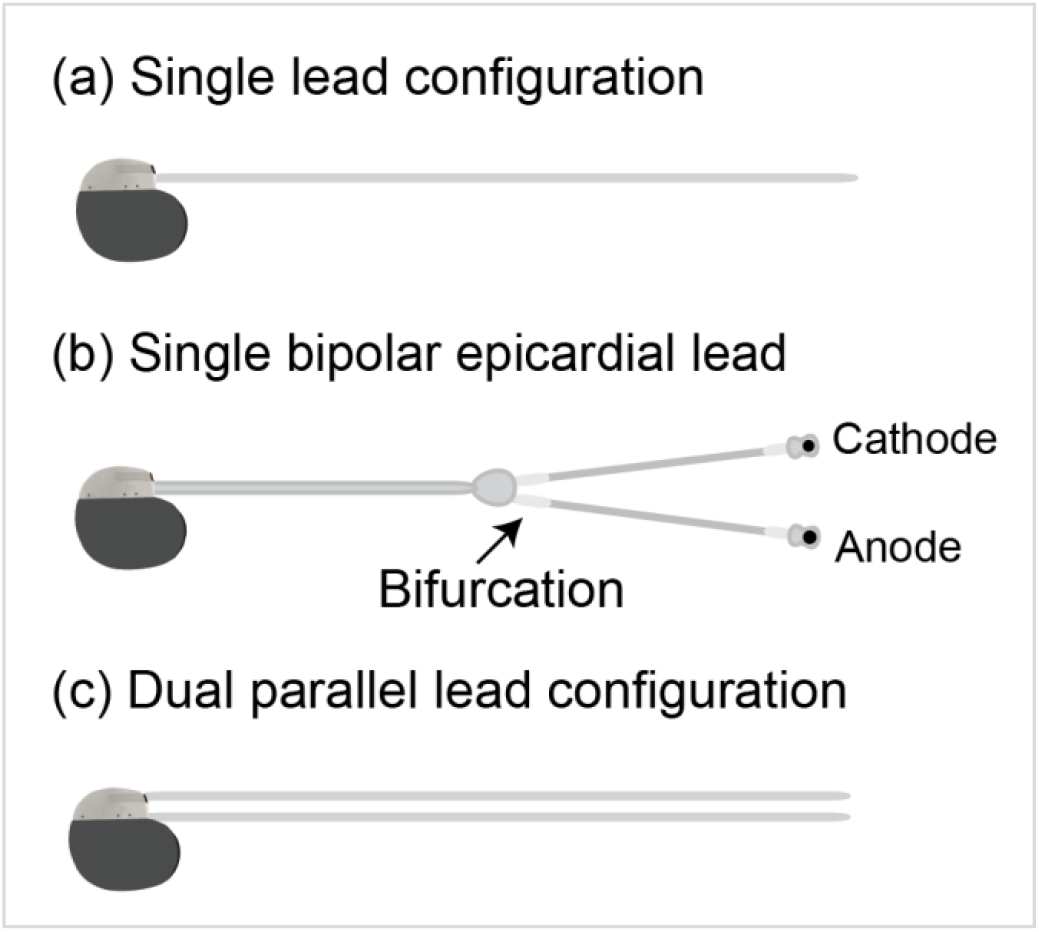
Comparison of lead configurations. (a) A simple, unbranched lead. (b) The bipolar bifurcated epicardial lead investigated in this study, in which a single lead body splits into two branches ending in separate electrodes. (c) A dual-lead system with two parallel, unconnected leads.

In this work, we introduce and experimentally validate a cumulative transfer function (cTF) framework to evaluate RF-induced heating for bifurcated leads. The method extends the conventional TF formalism by explicitly modeling branch-specific responses and cross-branch coupling, while reducing to the standard TF description in the limiting case of an unbranched geometry. We implement the framework for a commercial bipolar epicardial lead and evaluate its predictive performance against a simplified single-branch TF approximation that neglects branching effects. Validation is performed first in a homogeneous phantom across a set of canonical lead trajectories and then, without recalibration, in a heterogeneous anthropomorphic pediatric phantom using clinically derived trajectories. We further examine the sensitivity of the framework to the electrode separation used during cTF acquisition to determine whether predictions remain stable across modest variations in measurement geometry. Finally, we apply the validated cTF to adult and pediatric virtual human models to examine how predicted RF heating varies with anatomy, trajectory, and imaging landmark for the tested lead at 1.5 T. Through this framework, we aim to establish a validated method for quantifying RF-heating risk in bifurcated leads and to demonstrate how strongly that risk can depend on device geometry, anatomy, and exposure scenario.

## 2. Methods

Our workflow comprises four stages. Section 2.1 briefly recapitulates the established TF formalism to define the notation, lead domains (trunk and branches), and assumptions used in the cTF extension for bifurcated leads. Section 2.2 describes the reciprocity-based TF measurements and experimental setup used to obtain the cTFs. Section 2.3 presents estimation of the calibration (*α*) and coupling (*β*) parameters and validates the model first against experimental heating measurements for a set of canonical lead trajectories in an ISO/TS 10974 Tier-3 and ASTM-type phantom, and then in a heterogeneous anthropomorphic phantom supporting clinically derived, anatomically realistic trajectories. Finally, the validated cTF model is applied to predict in-vivo RF heating across virtual human models and imaging landmarks.

### 2.1 Theoretical Framework and Notation

We include a brief recap of the Tier-3 TF formalism solely to establish the notation, units, and assumptions used by our cTF generalization to bifurcated leads. We denote by *s* ∈ [0, *L*] the coordinate along the shared trunk, by *u* ∈ [0, *L*_*c*_] the coordinate along the cathode branch, and by *v* ∈ [0, *L*_*A*_] the coordinate along the anode branch (geometry in Fig. 2a). Let *E*_∥_(·) be the incident tangential electric field along the lead centerline obtained from full-wave simulations without the implant (ISO/TS 10974 Tier-3 assumption).

**Figure 2:**
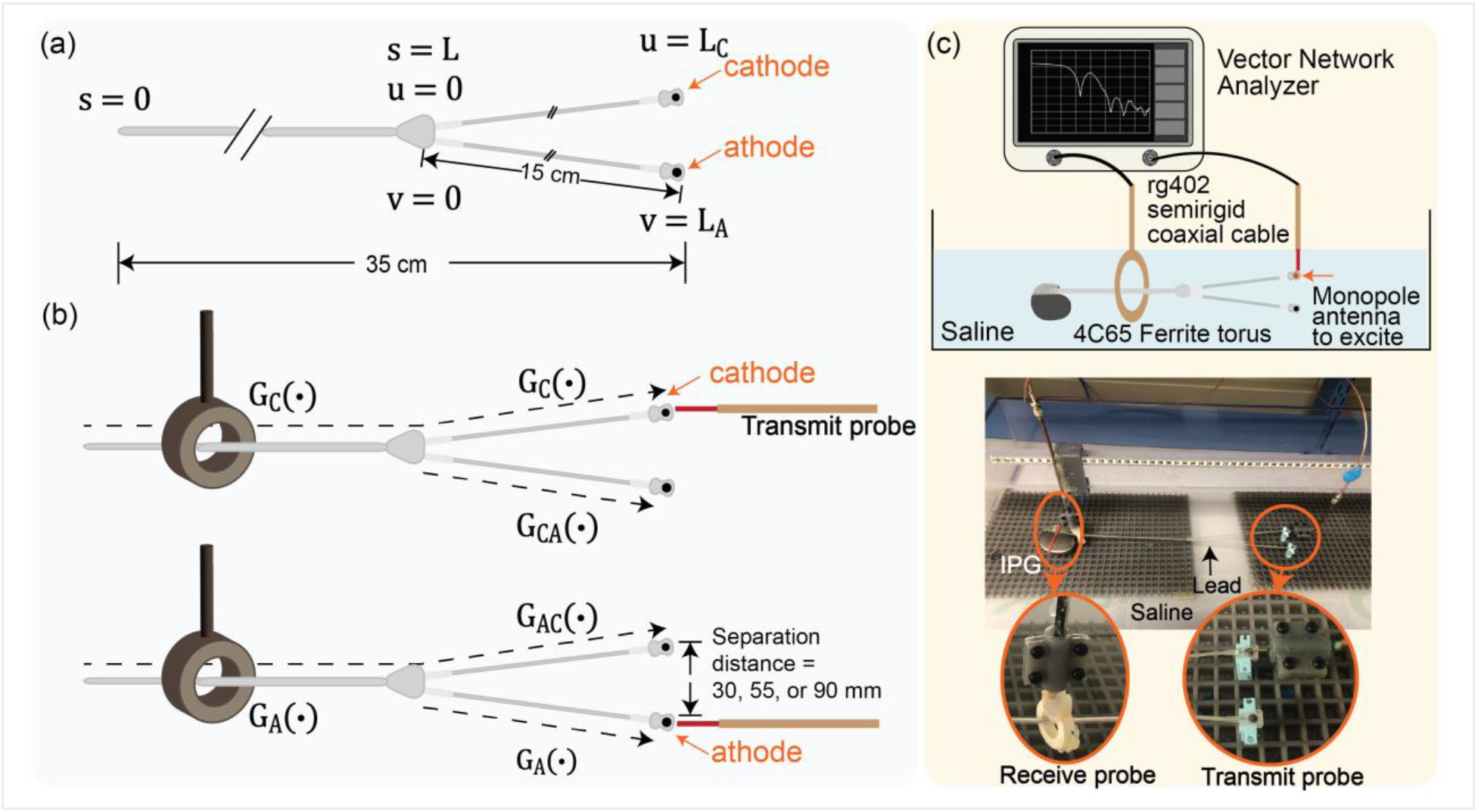
Experimental setup for cumulative transfer function (cTF) measurement. (a) Simplified rendering of the bipolar epicardial lead showing key dimensions and notations. (b) Schematic of the piecewise reciprocity-based measurement procedure used to obtain primary and cross-branch TF components, including the tested inter-electrode separation distance between the anode and cathode. (c) Measurement system including the VNA, transmit/receive probes, and the CIED submerged in the phantom.

For an unbranched lead, the heat-generating current induced around lead’s tip current is modeled as the line integral of the incident field weighted by a (generally complex) TF kernel *G*:

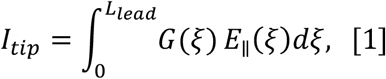

so that the temperature rise near the electrode obeys

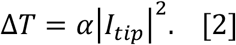

Here *α* > 0 is an empirically determined calibration constant (exposure-specific). Reciprocity-based measurements provide *G*(·) by exciting the electrode and recording the induced current along the lead (measurement schematic in Fig. 2b).

For a bifurcated geometry consisting of a trunk and two branches (cathode and anode), we model *self* and *cross-branch* responses explicitly. Defining separate TF kernels on each segment, the complex currents driving heating at the cathode (I_C_) and anode (I_A_) are:

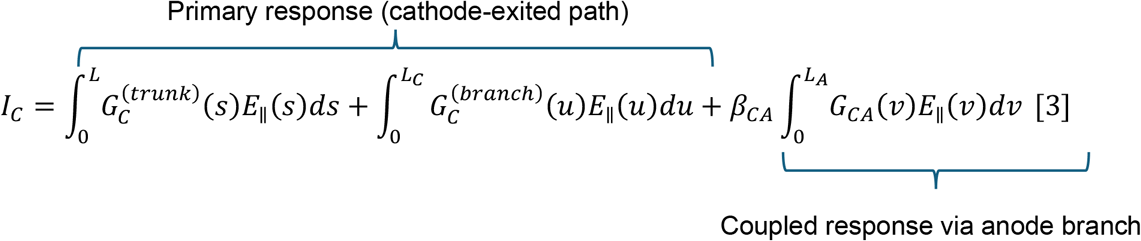

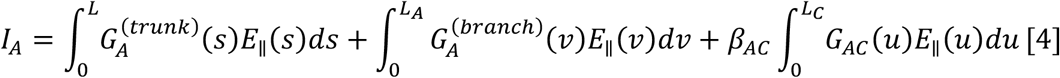

with corresponding temperature rises

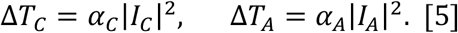

In Eqs. (3)-(5), 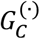 and 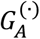 are primary TFs measured while exciting the cathode or anode, respectively, and *Gc*_*A*_, *G*_*A*_*c* are cross-branch TFs measured on the non-excited branch. *βc*_*A*_ and *β*_*A*_*c* are real-valued dimensionless coupling coefficients that account for junction-mediated effects not fully captured by the measured cross-branch kernels. *α*_*c*_ and *α*_*A*_ are electrode-specific calibration constants (real, >0) determined from phantom heating data. This formulation reduces to the conventional TF when there is no branching, and the cross-terms vanish (*βc*_*A*_ = *β*_*A*_*c* = 0).

We obtain the TF kernels in Eqs. (3)-(4) using the reciprocity setup of Fig. 2b: exciting one electrode and recording induced current along (i) the trunk and excited branch (primary TFs), and (ii) the non-excited branch (cross-branch TFs). Because inter-electrode spacing may influence coupling, we repeat these measurements at anode-to-cathode separations of 30, 55, and 90 mm and later show that prediction accuracy is insensitive to the separation used during TF acquisition. Additional experimental details follow in Section 2.2.

### 2.2 Transfer Function Measurement, Calibration, and Validation

All measurements were performed on a 35 cm bipolar epicardial pacing lead (CapSure® EPI 4968, Medtronic) connected to an Azure™ XT DR MRI SureScan IPG. This CIED was tested at 63.6 MHz (1.5 T).

#### 2.2.1 Reciprocity-based measurement of cumulative TFs (cTFs)

We measured the cTF kernels described in Eqs. (3)–(4) of Section 2.1 using a two-port VNA and a reciprocity setup (Fig. 2c),^10, 11, 13^ with the CIED submerged in saline (σ=0.5 S/m, ϵ_r_=80). A transmit probe excited one electrode (anode or cathode), while a receive probe was translated along the lead centerline to record the induced current proxy (complex *S*_21_) along (i) the trunk + excited branch (primary TFs 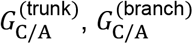) and (ii) the non-excited branch (cross-branch TFs *Gc*_*A*_, *G*_*A*_*c*). Sampling was performed every 10 mm along the shared trunk and every 5 mm along the branches. Measurements were repeated at inter-electrode separation distances of 30, 55, and 90 mm to probe coupling sensitivity to spacing. The complex *S*_21_ profiles (magnitude and phase) were used directly as uncalibrated TF kernels; absolute scaling is absorbed into the calibration constants *α*_*C*_, *α*_*A*_.

#### 2.2.2 Incident-field simulations and heating datasets for calibration and validation

For each experimentally evaluated lead routing, we first computed the incident tangential electric field *E*_∥_ along the trunk and each branch centerline using full-wave simulations that excluded the implant. Simulations were conducted in HFSS with a 1.5 T RF transmit body coil at 63.6 MHz, driven to 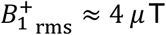, averaged over a 30-mm-radius circular plane centered at the coil’s calculate the incident tangential electric field (*E*_∥_) for a *B*^+^ ≈ 4 μ*T* on a transverse plane at isocenter. isocenter; complex *E*_∥_ values were then extracted along the candidate paths (Fig. 3). These fields were paired with the measured cumulative TF kernels and utilized in Eqs. (3)–(5) to generate heating predictions for comparison with temperature measurements.

**Figure 3:**
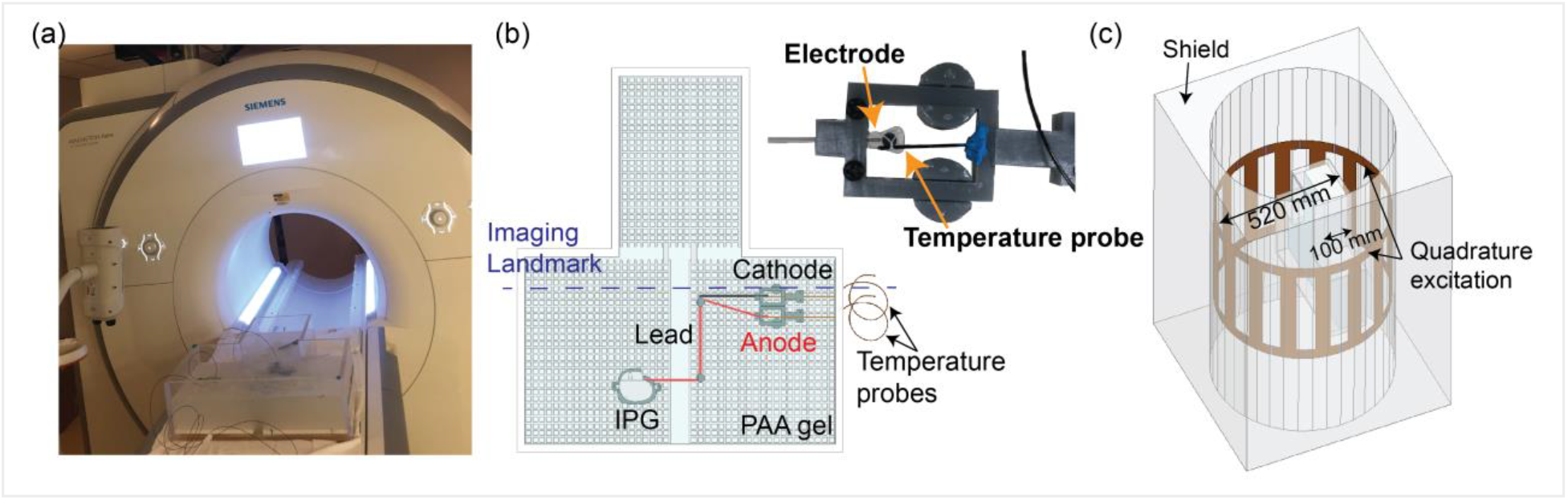
Calibration and validation workflow. (a) The experimental setup showing the CIED in the phantom inside the 1.5 T MRI scanner. (b) A 3D rendering of a validation trajectory within the phantom, with a fiber optic probe measuring temperature at an electrode. (c) Corresponding simulation setup, with the phantom model loaded into the RF body coil to calculate the incident tangential electric field (*E*_∥_) for a 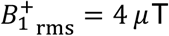on a transverse plane at isocenter.

Model calibration and standard validation used a canonical trajectory set acquired in a homogeneous polyacrylamide (PAA) gel phantom (σ=0.47 S/m, ε_r_=89.09). The CIED was routed along 24 configurations (Supplementary Fig. S1), and the temperature rise, Δ*T*, at both electrodes was measured with fiber-optic probes (Osensa, Burnaby, BC, Canada, 0.01– resolution) during a T1-TSE sequence running on a 1.5 T Siemens Aera scanner (TA = 280 s,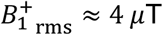).

To assess generalization under clinically relevant routings and heterogeneous media, we conducted a secondary validation in an anthropomorphic child-sized phantom with lead trajectories derived from clinical scans of patients with congenital heart disease and epicardial leads, using the same scanner and exposure conditions as in the canonical trajectory set (Supplementary Material Fig. S2). This step probes anatomy- and trajectory-dependent coupling not captured in a homogeneous gel phantom.

Electrode-specific calibration constants (*α*_*A*_, *α*_*C*_) and dimensionless coupling coefficients (*β*_*A*_*c, βc*_*A*_) were estimated from the canonical trajectory dataset using two separate joint optimizations that minimized the root-mean-square error between predicted and measured electrode temperature rise. For the cathode,

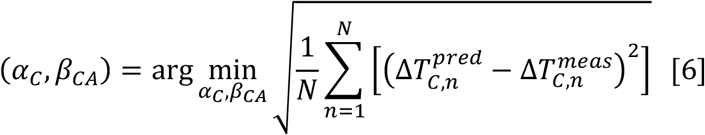

and (*α*_*A*_, *β*_*A*_*c*) were estimated analogously by minimizing the RMSE in Δ*T*_*A*_. Here, N is the number of trajectories (24 in this case). During optimization, *α*_*A*_and *α*_*c*_ were constrained to be positive; *β*_*A*_*c* and *βc*_*A*_ were estimated as real-valued unconstrained parameters.

The estimation was evaluated for 30, 55, and 90 mm inter-electrode separation distances, with the calibration constants and coupling coefficients fixed to those derived from the 30 mm cTFs in all cases, since the transmit probe location was kept the same for each separation distance. As a comparator, a single-branch TF model was fit for the cathode by setting *βc*_*A*_ = 0 and re-estimating *α*_C_ and vice versa for the anode. Performance metrics for both models and both datasets are reported in Section 3.2.

Note that the temperature rise scales with the squared transmit field amplitude:

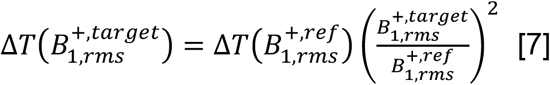

Here we report Δ*T* at a fixed RF exposure 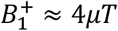, TA =280 s, which can be scaled to different RF conditions.

### 2.3 Prediction of in-vivo RF Heating

After calibration and validation, the cTF model was applied to predict in-vivo RF heating across a library of adult and pediatric virtual human body models (VHMs) (Fig. 4). Ten adult and ten pediatric VHMs were used; for each population, one model had nominal dielectric properties, and nine additional models were generated to represent inter-subject heterogeneity by independently scaling the conductivity and relative permittivity of each tissue by random factors drawn between 0.5 and 1.5 times the nominal values. The detailed dielectric property values of tissues used in the human body model simulations are reported in Supplementary Table S1.

**Figure 4:**
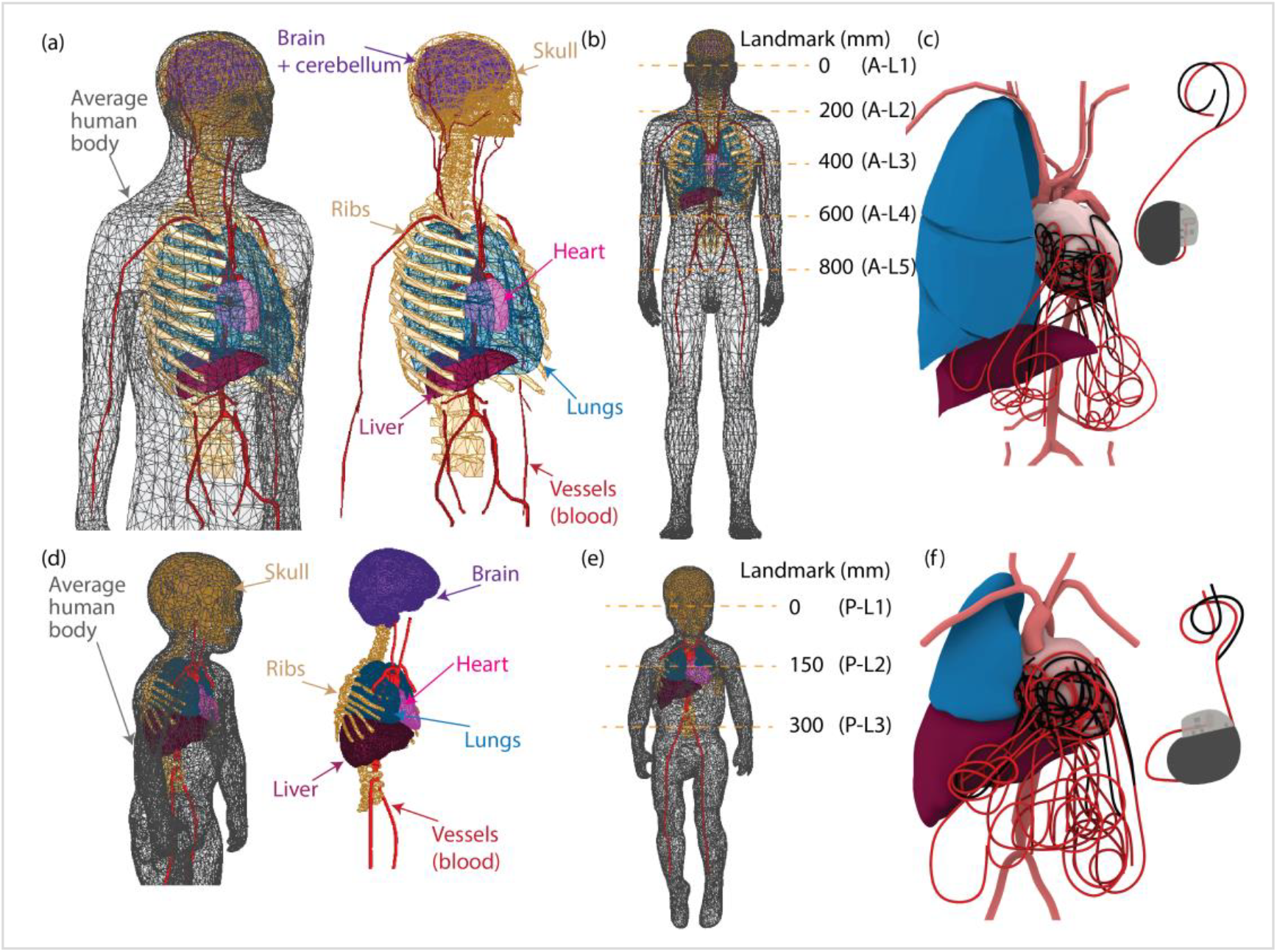
Setup for in-vivo RF heating predictions. (a) Example adult human body model and (d) pediatric body model, each comprising multiple tissue types. (b, e) Imaging landmarks evaluated in the adult and pediatric models, respectively. Each landmark is the position of the RF coil isocenter along the head-to-foot axis, given as the offset in millimeters from the nose (0 mm at the level of the nose, increasing toward the feet). The five adult landmarks are A-L1 (0 mm, head), A-L2 (200 mm, upper thorax), A-L3 (400 mm, thorax at heart level), A-L4 (600 mm, upper abdomen), and A-L5 (800 mm, lower abdomen and pelvis). The three pediatric landmarks are P-L1 (0 mm, head), P-L2 (150 mm, thorax), and P-L3 (300 mm, abdomen). (c, f) Library of 15 clinically relevant device trajectories evaluated in each body model.

For each VHM, fifteen clinically relevant device trajectories were evaluated (Fig. 4c,f). Predictions were computed at five imaging landmarks in the adult models (A-L1 to A-L5, head to pelvis) and three in the pediatric models (P-L1 to P-L3, head to abdomen); each landmark fixes the RF coil isocenter at a defined position along the head-to-foot axis (Fig. 4b,e). Incident tangential electric fields *E*_∥_ were obtained from full-wave simulations with the implant excluded (Tier-3 “no-implant” fields), using the same 1.5 T body coil model as in Section 2.2, driven to 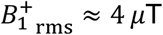. Complex *E*_∥_ values were extracted along the trunk and each branch centerline for every routing/landmark combination. (see Supplementary Material Fig. S3 for the simulation setup)

For each scenario, electrode currents *I*_C_ and *I*_*A*_ were computed via Eqs. (3)–(4) using the measured TF kernels and the coupling coefficients *β* estimated in Section 2.2; temperature rise at each electrode was then predicted from Eq. (5) using the electrode-specific calibration constants *α*_C_ and *α*_*A*_. Predictions were reported at the reference exposure used for calibration (TA = 280 s, 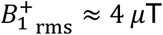), and then were scaled to estimate 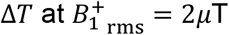.

To assess sensitivity to the electrode spacing used during TF measurement, the full set of predictions was generated separately for the three cTF sets acquired at inter-electrode separation distances of 30, 55, and 90 mm. In total, this resulted in 1,200 unique field scenarios (750 adult, 450 pediatric), each a body-model × trajectory × landmark combination. Evaluated at both electrodes and across the three cTF measurement separations, this produced 7,200 heating predictions (4,500 adult, 2,700 pediatric) at the 4 μT reference exposure, each then rescaled to 2 μT via Eq. 7.

### 2.4 Statistical Analysis

All analyses were performed in R (v4.1.2). For the primary (ASTM-type phantom) validation, predictive accuracy of the single TF and cTF models was quantified by mean absolute error (MAE), standard deviation of the error, error range, and root-mean-square error (RMSE), computed from the difference between predicted and measured Δ*T* at each electrode across the 24 canonical configurations.

For the in-vivo predictions, we first assessed normality of Δ*T* distributions using the Shapiro–Wilk test. If normally distributed, we used ANOVA; otherwise, we used the Kruskal–Wallis rank-sum test to evaluate whether the electrode separation distance used during TF measurement (30, 55, 90 mm) had a statistically significant effect on predicted Δ*T*. Tests were performed separately for adults and pediatrics and for anode and cathode predictions, aggregating across all VHMs, trajectories, and landmarks. All tests were two-sided with *α* = 0.05.

## 3. Results

### 3.1 Cumulative Transfer Function Measurements

We measured the magnitude and phase of the cTFs at electrode separation distances of 30, 55, and 90 mm, exciting each electrode in turn and sampling along the excited (primary) and non-excited (coupled) branches (Fig. 5). The data show asymmetry between anodal and cathodal excitation (both magnitude and phase) and a non-negligible coupled response on the non-excited branch, confirming that a single-TF is insufficient to predict heating at both electrodes. We observe no systemic, separation-dependent shift across the three TF-measurement separations beyond small local variations.

**Figure 5:**
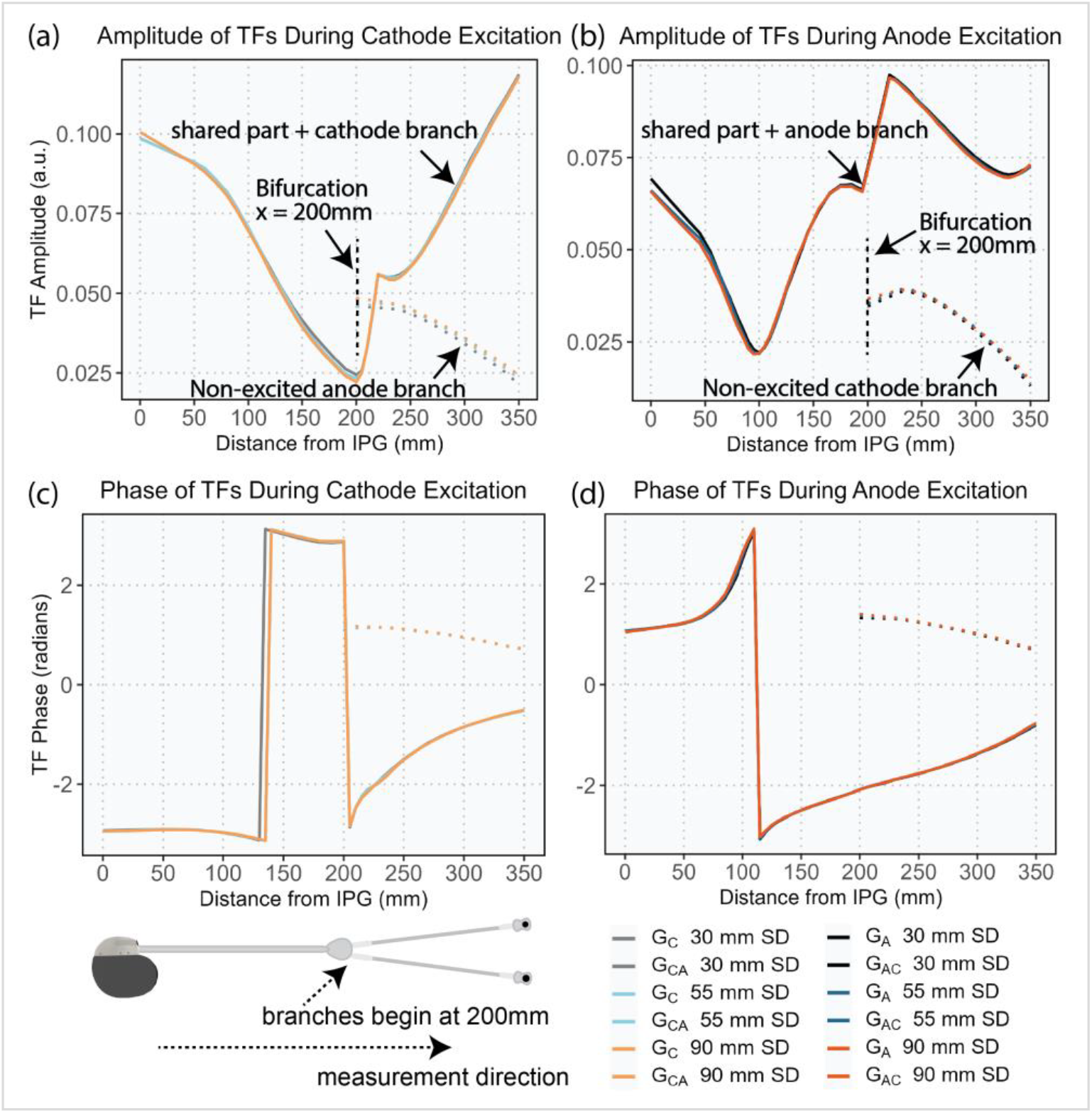
Measured cumulative transfer functions (cTFs) at three inter-electrode separation distances (SD = 30, 55, and 90 mm). Top panels show TF magnitude and bottom panels show phase. (a, c) TFs measured during cathode excitation, including the primary TF along the cathodal branch (G_C_) and the coupled TF along the anodal branch (G_CA_). (b, d) TFs measured during anode excitation including G_*A*_ and G_AC_. The data demonstrate clear asymmetry between cathodal and anodal excitation and confirm a non-negligible coupled response on the non-excited branch.

### 3.2 Cumulative Transfer Function Validation in ASTM-Type and Pediatric Phantoms

We validated both the single-TF comparator and the cTF model using 24 canonical lead trajectories routed in a homogeneous PAA-gel phantom (Fig. 6), spanning 35–245 mm inter-electrode separations (trajectory set in Supplementary Fig. S1). Scanner exposure was normalized to 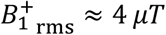. At each of the 24 configurations, Δ*T* was measured at both electrodes (N = 48 electrode-level observations).

**Figure 6:**
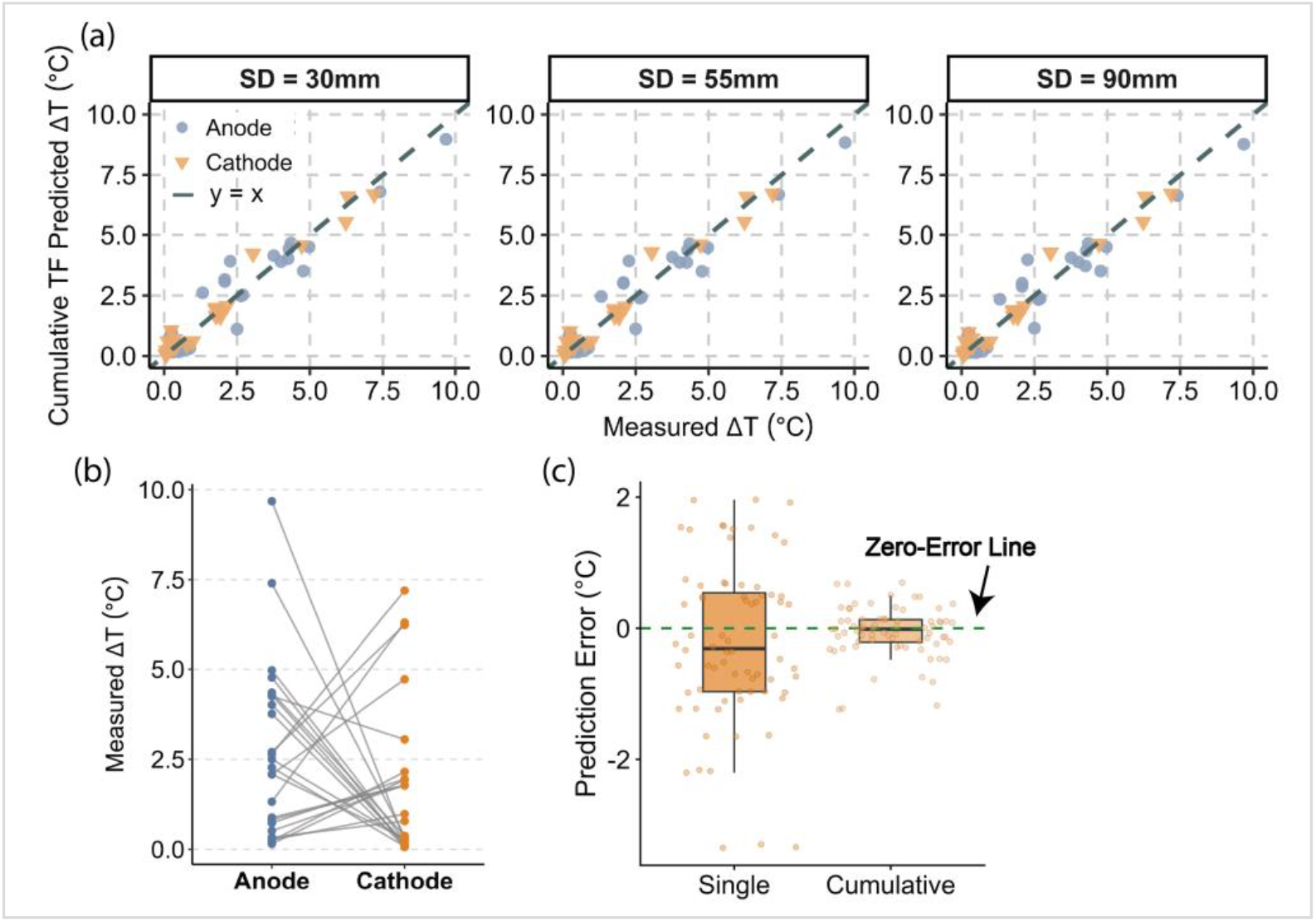
Validation of the cTF framework against experimental heating measurements for TF datasets acquired at three inter-electrode separation distances (30, 55, and 90 mm). (a) Scatter plots of measured versus predicted Δ*T* for anode and cathode electrodes; the dashed line denotes perfect agreement. (b) Measured Δ*T* at the anode versus cathode across the 24 validation configurations, demonstrating distinct thermal responses at the two electrodes. (c) Distribution of cathodal prediction error for the single-branch TF and cTF models, illustrating the improved accuracy and reduced error spread of the cTF approach. Cathodal error is shown because the single-branch approximation primarily degrades prediction at the cathode, whereas anodal error changed minimally between the two models.

Although Fig. 5 shows that the measured cTF curves (magnitude/phase) are visually similar across the three TF-measurement separation distances (30, 55, and 90 mm), small complex-valued differences could, in principle, propagate through Eqs. (3)–(5)—after integration along the trunk/branches and squaring—to affect Δ*T*predictions. To demonstrate prediction-stage robustness, we repeated the validation using each cTF dataset in turn, while applying the same calibration and coupling coefficients obtained from the 30 mm configuration. Fig. 6 reports the resulting accuracy separately for cTFs measured at 30, 55, and 90 mm. This separation-specific reporting complements Fig. 5’s measurement-stage invariance by confirming that prediction-stage accuracy is likewise insensitive to electrode separation during TF measurement.

Using the single-TF approach (measured along cathode branch, ignoring the non-excited anode branch), the mean absolute error (MAE) was 1.01 ± 0.74 °C, 1.01 ± 0.75 °C, and 1.00 ± 0.75 °C for TFs measured at 30, 55, and 90 mm electrode separation, respectively; the corresponding RMSE values were 1.24 °C, 1.25 °C, and 1.24 °C, with an error range of −3.35 to 1.96 °C (Fig. 6c). In contrast, the cTF reduced error across all three separations, with MAE = 0.29 ± 0.28 °C (30 mm), 0.28 ± 0.29 °C (55 mm), and 0.27 ± 0.29 °C (90 mm), and RMSE = 0.39–0.40 °C, narrowing the error range to −1.24 to 0.7 °C (Fig. 6c). Thus, cTF predictions achieve substantially higher fidelity and remain stable regardless of which TF-measurement separation is used to build the model.

As a complementary check of generalization beyond homogeneous, canonical routings, we performed a secondary validation in a heterogeneous anthropomorphic pediatric phantom with clinically derived trajectories (Supplementary Fig. S2). Without recalibration (using cTFs measured at 30 mm), the cTF reproduced measured heating with MAE = 0.17 ± 0.14 °C (Head), 0.26 ± 0.16 °C (Chest), and 0.30 ± 0.26 °C (Abdomen), with RMSE = 0.22–0.39 °C and error ranges from −0.92 °C to 0.67 °C. (Results pooled across both electrodes). Electrode-wise prediction performance and error statistics are reported in Supplementary Fig. S4 and Table S2, respectively, supporting robustness beyond the ASTM-type phantom.

*Note on terminology:* the separation distances reported here (30/55/90 mm) pertain to the spacing used during TF measurement in saline (Fig. 2b&c), whereas the 24 canonical validation trajectories in gel span 35–245 mm inter-electrode separations (Supplementary Fig. S1). Reporting both clarifies that the cTF’s predictive performance is insensitive to the TF-measurement separation even as actual trajectory separations vary widely in the validation set.

#### 3.3 Prediction of RF Heating in Virtual Human Models

The validated cTF was applied to 10 adult and 10 pediatric VHMs across 15 clinically relevant trajectories and multiple landmarks (five adult; three pediatric). This yielded 1,200 unique scenarios (body model × trajectory × landmark) and, accounting for two electrodes and three TF measurement separations, 7,200 predictions (4,500 adult; 2,700 pediatric) at an exposure level of 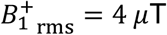 (and equivalently 7,200 normalized predictions at 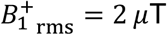).

In adults, worst-case predicted ΔT was strongly landmark-dependent (Fig. 7a; Table S3). Heating peaked at the heart and upper-abdominal landmarks, reaching 4.9 °C at A-L3 (anode) and 7.3 °C at A-L4 (cathode), the highest value across all adult models, trajectories, and electrodes. Worst-case ΔT fell to 1.3 °C at the pelvis (A-L5) and to 0.4 °C and 0.2 °C at the upper thorax (A-L2) and head (A-L1), the last two within the cTF validation error (RMSE < 1 °C). In pediatrics, heating was lower and again concentrated below the head, with worst-case ΔT of 2.0 °C at the chest (P-L2) and 2.2 °C at the abdomen (P-L3) versus 0.4 °C at the head (P-L1) (Fig. 7b; Table S4). For the 35 cm lead at 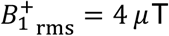, only the adult heart and upper-abdominal landmarks produced heating well into a potentially hazardous range, while the highest pediatric value was roughly one-third of the adult maximum. At 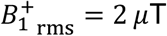, worst-case heating remained below 2 °C across all models and landmarks.

**Figure 7:**
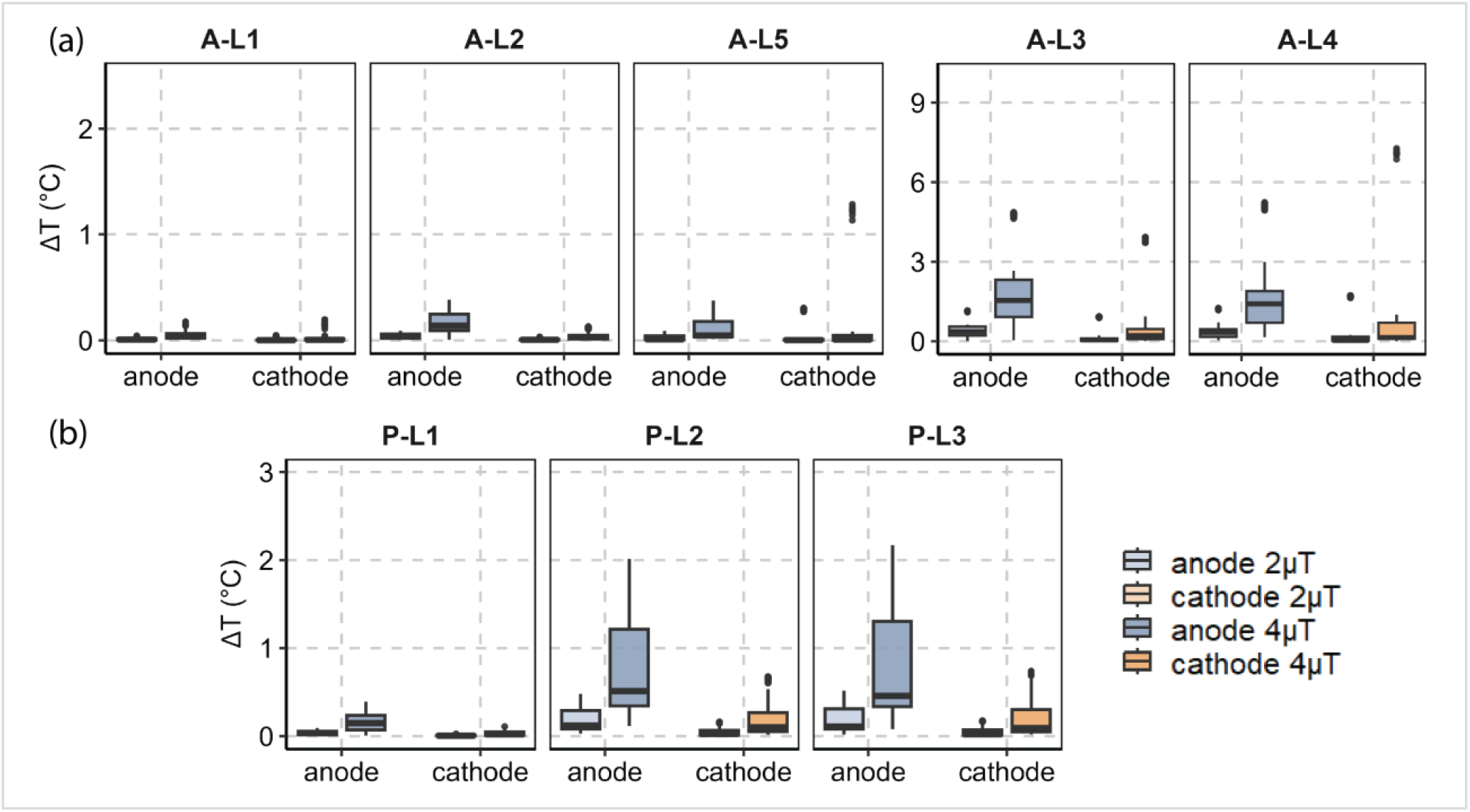
In-vivo ΔT predictions using the validated cTF for 10 adult (a) and 10 pediatric (b) body models, shown here for the cTF acquired at 30 mm inter-electrode separation. Panel (a) summarized results across five adult imaging landmarks, and panel (b) across three pediatric landmarks (see Fig. 4). Predictions are shown for 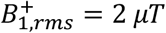 and 4 μ*T*with an exposure time of approximately 4 min 40 s.

Means differed by at most 0.02 °C across 30/55/90 mm TF-measurement separations for any landmark/electrode in Table S3 and S4, consistent with the Kruskal–Wallis tests showing no significant differences in adults (anode χ^2^(2)=0.078, p=0.962; cathode χ^2^(2)=0.071, p=0.965) or pediatrics (anode χ^2^(2)=0.318, p=0.853; cathode χ^2^(2)=0.006, p=0.997). Together, these indicate robustness of the cTF predictions to the electrode separation used during TF acquisition.

## 4. Discussion

Assessing RF heating of implants with elongated leads is a substantial challenge, as its magnitude is a complex interplay of the implant’s geometry and material, its trajectory within the body, the patient’s size and composition, and the specific MRI scanner.^18-24^ While standardized assessment methods have enabled MRI-conditional labeling for many devices with simple, unbranched leads, implants with bifurcated leads remain a significant hurdle.^13, 14, 25-29^ We introduced and experimentally validated a cumulative TF framework that explicitly accounts for branch-specific response and inter-branch coupling in bifurcated leads. Relative to a single-branch TF approximation, the cTF reduced prediction error for the tested 35 cm bipolar epicardial lead by nearly 70% (RMSE ≈1.24 °C → 0.4°C; MAE ≈1°C → 0.3 °C), narrowing error ranges (Fig. 6). These improvements held regardless of which cTF set—measured at 30, 55, or 90 mm electrode separation—was used to make predictions.

### 4.1 Why the cTF Helps: Explicit Treatment of Asymmetry and Coupling

Two empirical observations motivate the cTF. First, anode and cathode responses are asymmetric in both magnitude and phase, so a single TF cannot serve both ends of a bifurcated device (Fig. 5). Second, we measured non-negligible induced current on the non-excited branch under tip excitation, directly evidencing inter-branch coupling. In that setting, a single-lead model that assumes the other branch is electrically invisible will mis-estimate the superposed current that ultimately drives heating. By separating primary and cross-branch responses and allowing for a coupling coefficient, the cTF better captures the actual current delivered to each electrode under distributed MRI fields, which is reflected in the lower prediction errors (Fig. 6).

### 4.2 Clinical Implications and Practical Workflow

Epicardial pacing systems are essential for pediatric patients and adults for whom transvenous implantation is not feasible. However, the lack of MR-conditional labeling for these devices, driven by the assessment challenges of bifurcated leads, often forces these patients to undergo alternative imaging modalities often involving ionizing radiation. Our group has previously shown that for simpler lead geometries, strategically modifying the implant’s trajectory can substantially reduce its coupling with MRI electric fields and thus minimize RF heating.^20, 30, 31^ The validated cTF model presented here provides the critical framework needed to apply this same promising optimization approach to bifurcated leads, creating a pathway to identify safer implantation trajectories and expand MRI access.

Our in-vivo predictions using this new model offer immediate insights into clinical risk and show that these risks cannot be generalized across patient populations. For the pediatric models, the predicted temperature rise remained low across all trajectories and landmarks (max ΔT of 2.2°C), indicating that for the tested 35 cm lead at 1.5 T, predicted RF heating remained low across the evaluated pediatric anatomies, trajectories, and imaging landmarks. However, this finding is specific to the 35 cm lead evaluated and should be interpreted with caution. Lead length is a critical determinant of resonant behavior, and these results should not be extrapolated. Indeed, a previous study using direct phantom measurements found that a shorter, 25 cm epicardial lead produced a maximum temperature rise of ∼12°C in a pediatric model—substantially higher than the heating observed from the 35 cm lead under similar conditions.^31^ This highlights the necessity of specific assessment for each device model. In contrast, the adult models showed substantially higher predicted heating in selected chest and upper abdominal scenarios, with maximum predicted Δ*T*= 7.3°C. Taken together, these findings illustrate that RF-heating risk in bifurcated epicardial systems is highly scenario-dependent and should not be inferred from a single anatomy, routing, or imaging landmark.

#### 4.3. The Effect of TF-Measurement Separation Distance and Robustness

Our investigation into the effect of electrode separation distance during the initial TF measurement revealed that this parameter had a minimal impact on the final heating predictions. While we observed minor variations in the raw TF measurements at different distances, these differences did not translate into statistically significant changes in the predicted in-vivo temperature rise. This finding suggests that the cTF method is robust to small variations in the relative position of the electrodes, which adds confidence to its application in clinical scenarios where the exact in-vivo separation distance is not known.

## 5. Conclusion

Our results reveal that RF heating risks are highly dependent on patient anatomy, with pediatric models showing a low risk of significant heating for the specific lead length tested, while adult models exhibited potentially hazardous temperature increases at abdominal imaging landmarks. It is critical to note that these findings are specific to the 35 cm lead model evaluated; RF heating behavior can change significantly with lead length, and these results should not be generalized to other epicardial lead models without specific assessment. The framework presented here provides a robust and validated pathway for the safety assessment of a wide range of complex implantable devices with bifurcated leads, which is a critical step toward enabling MR-conditional labeling and expanding safe MRI access for all patients.

