## Supplementary Material for "Cumulative Transfer Function for Assessment of MRI-Induced RF Heating Risk in Pediatric Patients Implanted with Bifurcated Leads"

**Supplemental**

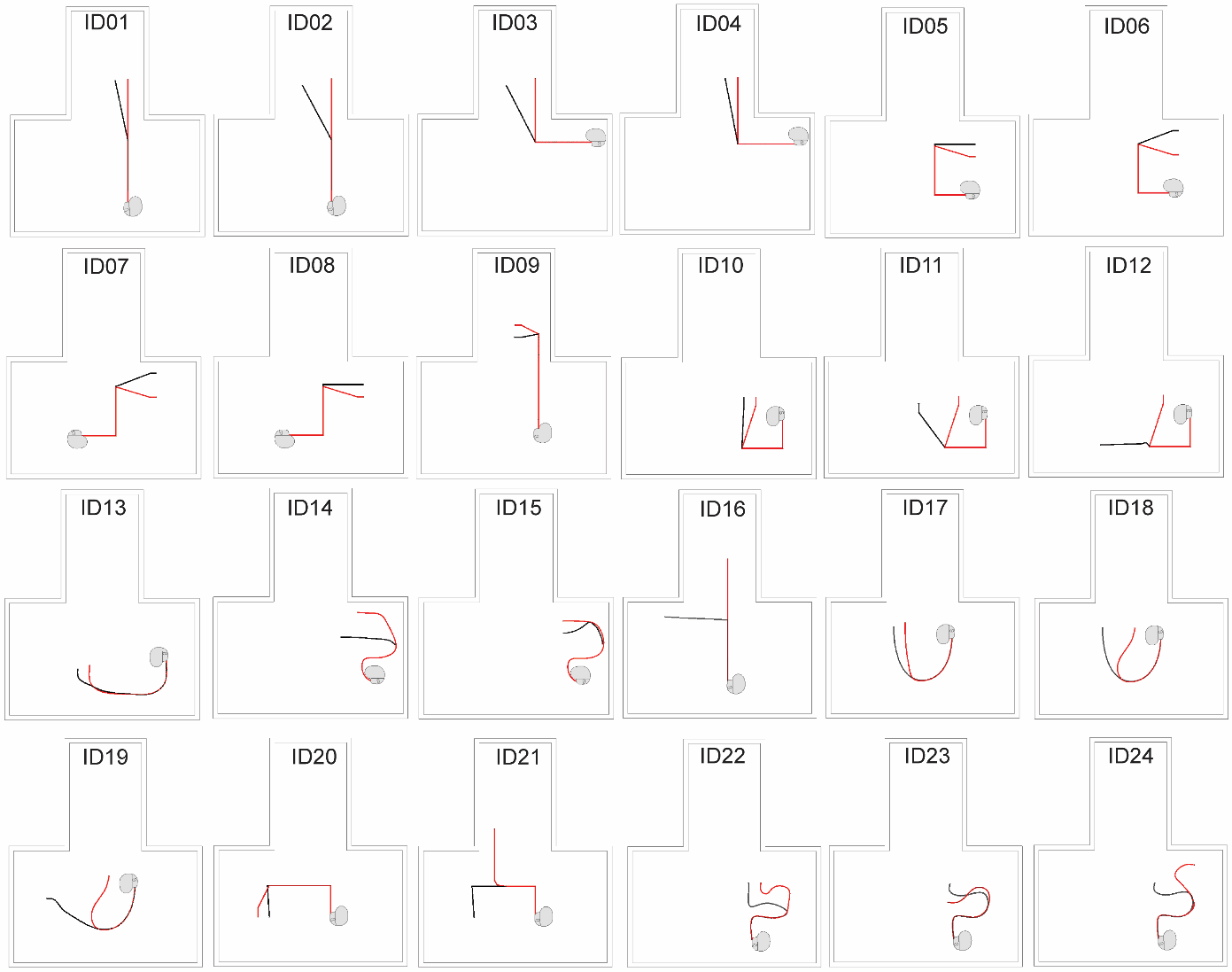

**Figure S1:** 24 canonical lead trajectories used for calibration and primary validation of the cTF framework in the ASTM-type phantom. The anodal branch is shown in red while the cathodal branch is shown in black. The shared proximal trunk is shown in red for visualization clarity. The separation distance between the anode and cathode ranges from 35-245 mm.

**Validation of Cumulative Transfer Function (cTF) with Clinically Relevant Bipolar Epicardial Lead Trajectories**

After calibration and validation in an ASTM-type homogeneous phantom, the cTFs were applied without further recalibration in an anthropomorphic heterogeneous phantom to assess whether the calibrated cTF framework generalizes accurately beyond the calibration conditions.

To do this, we performed additional experiments with an anthropomorphic phantom constructed from segmentation of MRI images of a 29-month-old child. This phantom consisted of an agar-based heart structure (σ=0.68 S/m, 𝜀_r_=80.76) and polyacrylamide (PAA) gel (σ=0.47 S/m, 𝜀_r_ =89.09) that filled the remainder of the phantom shell. Within the phantom, we evaluated 10 unique, clinically relevant CIED configurations (Figure S2). These trajectories were created based on a retrospective analysis of radiographs of patients with epicardial pacemaker systems, with some directly replicated and others inspired by them to capture a broad range of temperature rise scenarios. Retrospective use of the imaging data was approved by Lurie Children’s Hospital of Chicago institutional review board. 3D-printed guides were used to shape the commercial lead for each trajectory. As in the primary validation done in the ASTM-type phantom, fiber-optic temperature probes were attached to the electrode to measure ΔT during RF exposure from the same T1-TSE sequence. The phantom was placed on the patient table in the head-first, supine position. Imaging was performed with the regions corresponding to the head, chest, and abdomen at the isocenter of the scanner.

We also simulated the distribution of the incident electric field inside the pediatric-sized phantom due to the RF transmit coil in the 1.5 T Siemens Aera system. The simulation parameters matched the experimental setup, and complex tangential E-field (Etan) values were extracted along each trajectory as previously described. RF heating was then predicted using the calibrated cTFs (measured with a 30 mm electrode separation).

Figure S4 presents the validation results, comparing measured and predicted ΔT values across three imaging landmarks. ${B_{1}^{+}}_{\mathrm{rms}}$ was normalized to approximately $4\text{ }\mu T$. Table S2 reports detailed error statistics for each electrode, including the mean absolute error ± standard deviation, RMSE, and error range. These results demonstrate that the cTF approach remains valid across clinically relevant imaging landmarks without requiring recalibration.

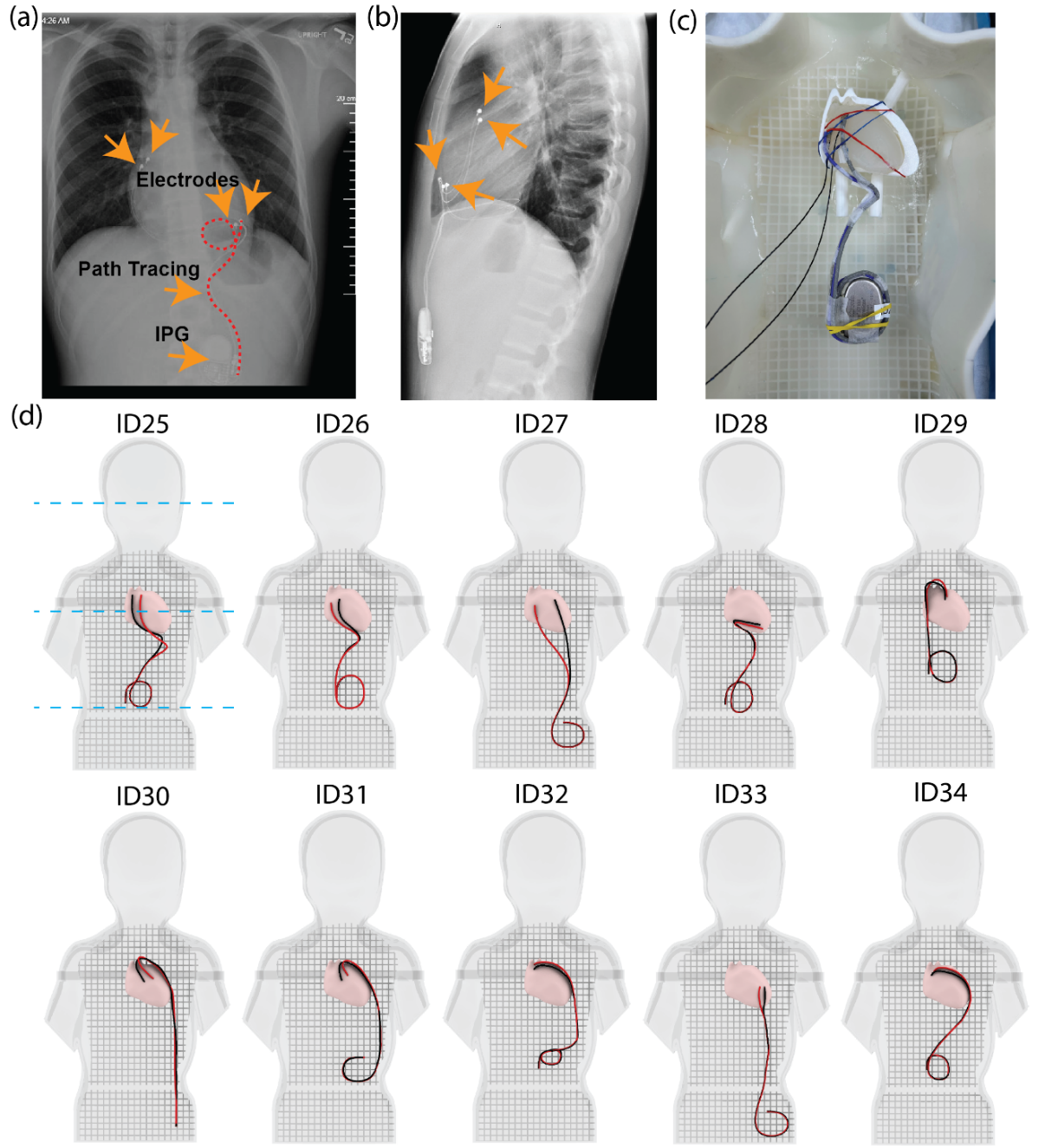

**Figure S2:** (a, b) Posterior-anterior (a) and lateral (b) chest radiographs from an adolescent patient, showing typical in-vivo pathways of implanted bipolar epicardial leads. (c) The experimental setup, where a 3D-printed guide was used to shape the lead into a specific, clinically relevant pathway. (d) An overview of the 10 trajectories evaluated within the anthropomorphic pediatric phantom. Blue dashed lines indicate the imaging landmarks (head, chest, and abdomen). For each trajectory, red lines represent the anodal branches, and black lines represent the cathodal branches.

**Table S1:** **Dielectric property values of tissues in human body models during simulations**

|  | **Body 1** | | **Body 2** | | **Body 3** | | **Body 4** | | **Body 5** | |
| --- | --- | --- | --- | --- | --- | --- | --- | --- | --- | --- |
| Tissues | ε**_r_** | σ (S/m) | ε**_r_** | σ | ε**_r_** | σ | ε**_r_** | σ | ε**_r_** | σ |
| Blood | 86.40 | 1.20 | 82.94 | 1.15 | 105.41 | 1.46 | 66.53 | 0.92 | 99.36 | 1.38 |
| Brain (average) | 82.60 | 0.40 | 90.86 | 0.44 | 106.55 | 0.52 | 56.99 | 0.28 | 89.21 | 0.43 |
| Cancellous bone | 30.90 | 0.20 | 25.34 | 0.16 | 34.30 | 0.22 | 18.23 | 0.12 | 44.81 | 0.29 |
| Cerebellum | 116.30 | 0.70 | 103.51 | 0.62 | 112.81 | 0.68 | 144.21 | 0.87 | 138.40 | 0.83 |
| Cortical bone | 16.60 | 0.06 | 23.57 | 0.09 | 8.80 | 0.03 | 12.28 | 0.04 | 17.43 | 0.06 |
| Heart | 106.50 | 0.70 | 53.25 | 0.35 | 71.36 | 0.47 | 151.23 | 0.99 | 97.98 | 0.64 |
| Liver | 80.60 | 0.50 | 74.15 | 0.46 | 78.18 | 0.49 | 102.36 | 0.64 | 92.69 | 0.58 |
| Lungs | 37.10 | 0.30 | 35.62 | 0.29 | 24.12 | 0.20 | 25.60 | 0.21 | 43.78 | 0.35 |
| Average human body | 78.00 | 0.47 | 99.06 | 0.60 | 65.52 | 0.39 | 61.62 | 0.37 | 88.92 | 0.54 |

|  | **Body 6** | | **Body 7** | | **Body 8** | | **Body 9** | | **Body 10** | |
| --- | --- | --- | --- | --- | --- | --- | --- | --- | --- | --- |
| Tissues | ε**_r_** | σ (S/m) | ε**_r_** | σ | ε**_r_** | σ | ε**_r_** | σ | ε**_r_** | σ |
| Blood | 95.90 | 1.33 | 115.78 | 1.61 | 124.42 | 1.73 | 129.60 | 1.80 | 97.63 | 1.36 |
| Brain (average) | 58.65 | 0.28 | 70.21 | 0.34 | 113.16 | 0.55 | 94.99 | 0.46 | 74.34 | 0.36 |
| Cancellous bone | 39.24 | 0.25 | 32.14 | 0.21 | 32.45 | 0.21 | 17.30 | 0.11 | 21.32 | 0.14 |
| Cerebellum | 140.72 | 0.85 | 134.91 | 0.81 | 88.39 | 0.53 | 120.95 | 0.73 | 110.49 | 0.67 |
| Cortical bone | 12.12 | 0.04 | 15.27 | 0.06 | 13.61 | 0.05 | 20.25 | 0.07 | 14.28 | 0.05 |
| Heart | 66.03 | 0.43 | 143.78 | 0.95 | 66.03 | 0.43 | 108.63 | 0.71 | 135.26 | 0.89 |
| Liver | 76.57 | 0.48 | 60.45 | 0.38 | 92.69 | 0.58 | 58.03 | 0.36 | 102.36 | 0.64 |
| Lungs | 35.62 | 0.29 | 41.18 | 0.33 | 36.36 | 0.29 | 22.26 | 0.18 | 53.42 | 0.43 |
| Average human body | 90.48 | 0.55 | 84.24 | 0.51 | 88.92 | 0.54 | 47.58 | 0.29 | 115.44 | 0.70 |

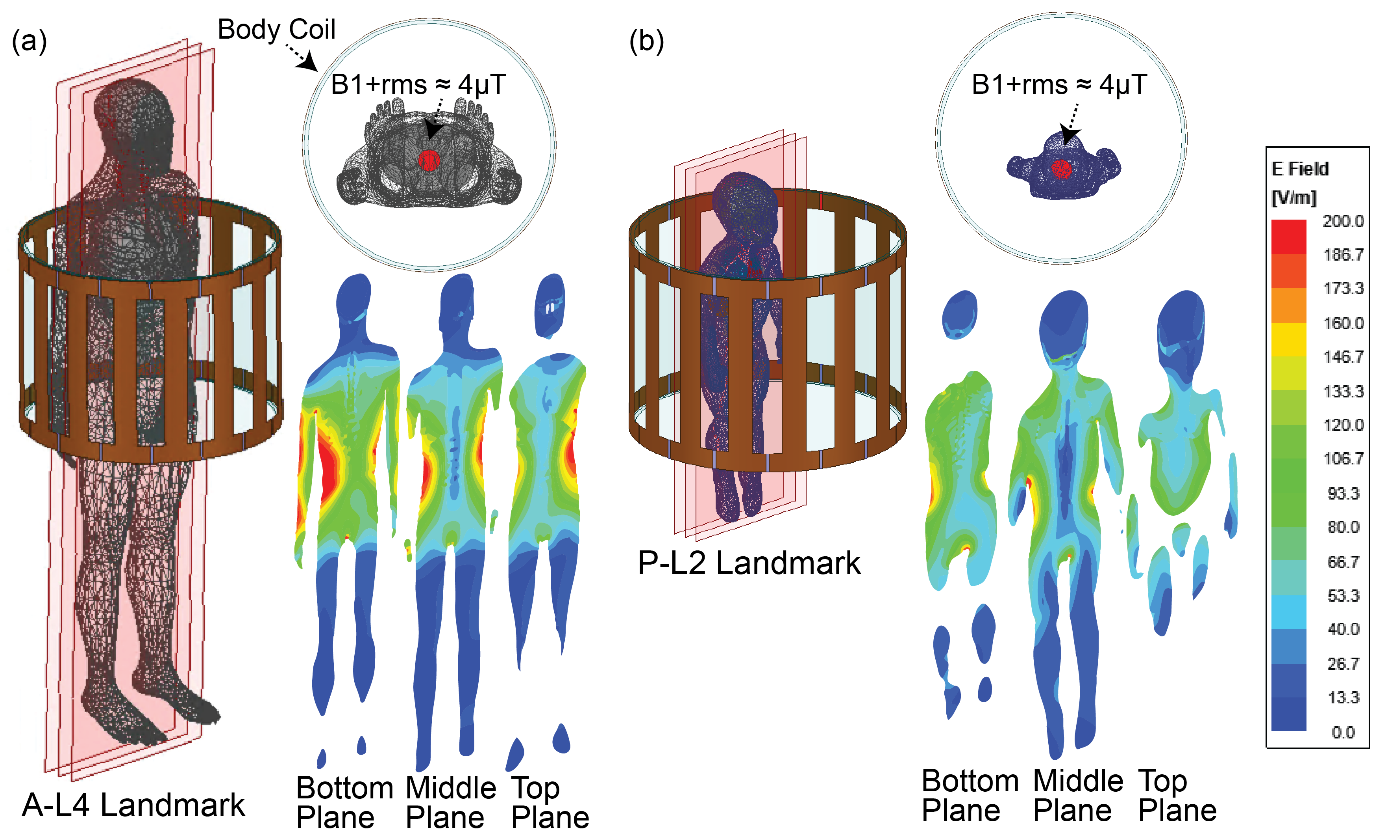

**Figure S3:** Simulation setup and E-field distribution for in-vivo RF heating predictions. The setups for predicting RF heating are shown for (a) the adult and (b) the pediatric virtual human models. For each, incident E-field distributions are displayed on three coronal planes, using models with nominal tissue properties. The examples shown are for the A-L4 landmark (adult) and the P-L2 landmark (pediatric). The RF exposure was normalized to ${B_{1}^{+}}_{\mathrm{rms}}\approx4\text{ }\mu T$, averaged over a circular plane of radius 30 mm centered within the model at the axial level of isocenter.

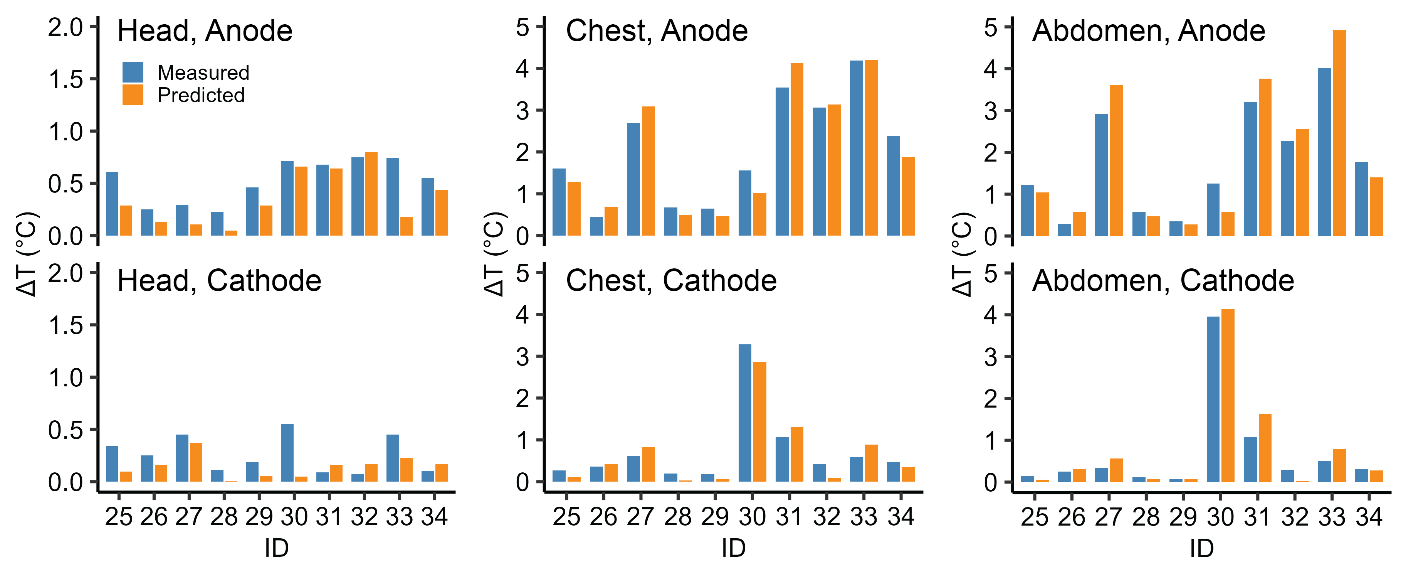

**Figure S4:** Secondary validation of the cTF framework in the anthropomorphic pediatric phantom. Measured and predicted temperature rise (ΔT) are compared for 10 clinically relevant device configurations across three imaging landmarks (head, chest, abdomen), using cTFs measured at a 30 mm electrode separation and applied without recalibration. RF exposure was normalized to ${B_{1}^{+}}_{\mathrm{rms}}\approx4\text{ }\mu T$.

**Table S2: Electrode-Wise Error Statistics for cTF ΔT Prediction in the Secondary Validation at** ${\boldsymbol{B}_{\boldsymbol{1}}^{\boldsymbol{+}}}_{\mathbf{rms}}\boldsymbol{\approx4}\text{ }\boldsymbol{\mu T}$

| Landmarks | Electrodes | MAE ± SD(\|error\|) (°C) | Error Range(°C) | RMSE(°C) |
| --- | --- | --- | --- | --- |
| P-L1 (Head) | Anode | 0.18 ± 0.16 | [-0.05, 0.56] | 0.23 |
| P-L1 (Head) | Cathode | 0.16 ± 0.13 | [-0.10, 0.50] | 0.20 |
| P-L2 (Chest) | Anode | 0.30 ± 0.20 | [-0.59, 0.54] | 0.36 |
| P-L2 (Chest) | Cathode | 0.21 ± 0.11 | [-0.30, 0.41] | 0.24 |
| P-L3 (Abdomen) | Anode | 0.41 ± 0.28 | [-0.92, 0.67] | 0.49 |
| P-L3 (Abdomen) | Cathode | 0.18 ± 0.17 | [-0.56, 0.26] | 0.24 |

**Table S3: Predicted ΔT Heating at** ${\boldsymbol{B}_{\boldsymbol{1}}^{\boldsymbol{+}}}_{\mathbf{rms}}\boldsymbol{\approx4}\text{ }\boldsymbol{\mu T}$ **in Adult Body Models Across Five Landmarks (Mean ± SD and Range)**

| TF Separation Distance | Adult Landmark | Anode heating (mean ± std) (°C) | Anode heating range (°C) | Cathode heating (mean ± std) (°C) | Cathode heating range (°C) |
| --- | --- | --- | --- | --- | --- |
| 30mm | A-L1 | 0.05 ± 0.04 | [0, 0.17] | 0.02 ± 0.04 | [0, 0.20] |
| 55mm | A-L1 | 0.05 ± 0.04 | [0.01, 0.17] | 0.02 ± 0.04 | [0, 0.20] |
| 90mm | A-L1 | 0.05 ± 0.04 | [0.01, 0.17] | 0.02 ± 0.04 | [0, 0.20] |
| 30mm | A-L2 | 0.16 ± 0.10 | [0.01, 0.38] | 0.04 ± 0.03 | [0, 0.13] |
| 55mm | A-L2 | 0.16 ± 0.10 | [0.01, 0.38] | 0.04 ± 0.04 | [0, 0.13] |
| 90mm | A-L2 | 0.16 ± 0.10 | [0.01, 0.38] | 0.04 ± 0.04 | [0, 0.13] |
| 30mm | A-L3 | 1.67 ± 1.15 | [0.03, 4.87] | 0.50 ± 0.93 | [0.01, 3.93] |
| 55mm | A-L3 | 1.65 ± 1.13 | [0.02, 4.79] | 0.50 ± 0.93 | [0.01, 3.94] |
| 90mm | A-L3 | 1.65 ± 1.13 | [0.02, 4.79] | 0.49 ± 0.94 | [0.01, 3.98] |
| 30mm | A-L4 | 1.57 ± 1.20 | [0.16, 5.24] | 0.73 ± 1.73 | [0, 7.27] |
| 55mm | A-L4 | 1.55 ± 1.19 | [0.14, 5.16] | 0.73 ± 1.73 | [0, 7.27] |
| 90mm | A-L4 | 1.55 ± 1.19 | [0.13, 5.17] | 0.73 ± 1.74 | [0, 7.29] |
| 30mm | A-L5 | 0.10 ± 0.11 | [0.01, 0.38] | 0.10 ± 0.30 | [0, 1.28] |
| 55mm | A-L5 | 0.10 ± 0.10 | [0.01, 0.35] | 0.10 ± 0.30 | [0, 1.28] |
| 90mm | A-L5 | 0.10 ± 0.10 | [0.01, 0.33] | 0.10 ± 0.30 | [0, 1.28] |

**Table S4: Predicted ΔT Heating at** ${\boldsymbol{B}_{\boldsymbol{1}}^{\boldsymbol{+}}}_{\mathbf{rms}}\boldsymbol{\approx4}\text{ }\boldsymbol{\mu T}$ **in Pediatric Body Models Across Three Landmarks (Mean ± SD and Range)**

| TF Separation Distance | Pediatric Landmark | Anode heating (mean ± std) (°C) | Anode heating range (°C) | Cathode heating (mean ± std) (°C) | Cathode heating range (°C) |
| --- | --- | --- | --- | --- | --- |
| 30mm | P-L1 | 0.15 ± 0.10 | [0.01, 0.39] | 0.03 ± 0.03 | [0, 0.11] |
| 55mm | P-L1 | 0.15 ± 0.09 | [0.01, 0.38] | 0.03 ± 0.03 | [0, 0.11] |
| 90mm | P-L1 | 0.15 ± 0.09 | [0.01, 0.38] | 0.03 ± 0.03 | [0, 0.10] |
| 30mm | P-L2 | 0.80 ± 0.58 | [0.11, 2.01] | 0.17 ± 0.18 | [0.02, 0.68] |
| 55mm | P-L2 | 0.79 ± 0.57 | [0.11, 1.96] | 0.17 ± 0.17 | [0.02, 0.67] |
| 90mm | P-L2 | 0.79 ± 0.56 | [0.11, 1.94] | 0.17 ± 0.17 | [0.02, 0.66] |
| 30mm | P-L3 | 0.80 ± 0.62 | [0.08, 2.17] | 0.19 ± 0.19 | [0.02, 0.74] |
| 55mm | P-L3 | 0.78 ± 0.61 | [0.07, 2.11] | 0.19 ± 0.19 | [0.02, 0.73] |
| 90mm | P-L3 | 0.78 ± 0.61 | [0.07, 2.10] | 0.19 ± 0.19 | [0.02, 0.72] |

Tables S3 and S4 show that predicted heating summaries remained highly stable across cTF datasets measured at 30, 55, and 90 mm separation, with minimal variation in the mean and range within each landmark.
